# A naïve piRNA Surveillance System That Broadly Monitors the Germline Transcriptome for Adaptive Genome Defense

**DOI:** 10.1101/2024.01.24.577019

**Authors:** Keisuke Shoji, Yukihide Tomari

**Affiliations:** Laboratory of RNA Function, Institute for Quantitative Biosciences, The University of Tokyo, 1-1-1 Yayoi, Bunkyo-ku, Tokyo, Japan; Department of Computational Biology and Medical Sciences, Graduate School of Frontier Sciences, The University of Tokyo, Bunkyo-ku, Tokyo, Japan; Graduate school of Bio-Applications and Systems Engineering, Tokyo University of Agriculture and Technology, Koganei-shi, Tokyo, Japan

## Abstract

PIWI-interacting RNAs (piRNAs) safeguard germline genomes from invasive genetic elements such as transposons. While long-term silencing relies on sequence-specific mechanisms such as ping-pong amplification and the incorporation of invader fragments into genomic piRNA clusters, how newly invading elements are initially recognized as “non-self” remains unresolved. Here, we show that low levels of sense-stranded piRNAs are broadly generated from the transcriptome in silkworms, flies, and mice, in direct proportion to RNA abundance and largely independent of canonical piRNA biogenesis pathways. This process can directly sample invaders: in silkworm cells, extremely abundant Bombyx mori latent virus RNAs enter this pathway. Reanalysis of recently endogenized retroviruses reveals distinct stages of adaptation in vivo—mouse AKV remains abundance-coupled, whereas koala KoRV-A has progressed into ping-pong amplification. We propose that this inefficient, non-selective, abundance-coupled pathway constitutes a naïve germline surveillance system that seeds an initial piRNA pool for later recruitment into more efficient, specificity-conferring silencing pathways.

## Introduction

PIWI-interacting RNAs (piRNAs) are a class of small RNAs of ∼24–32 nucleotides (nt) in length expressed in the germline. They associate with PIWI-subfamily proteins and guide them to repress invasive genetic elements such as transposons^1,2^. Many piRNAs are derived from genomic regions called piRNA clusters, which serve as reservoirs for transposon fragments and are often transcribed in the antisense orientation relative to transposon sequences^3,4^. Consequently, these piRNAs possess sequences complementary to transposons, allowing the piRNA-PIWI complex to recognize and cleave transposon mRNAs for silencing^5–8^. After the cleavage, the 3’ fragment of the cleavage product is subsequently loaded into another PIWI protein and acts as a precursor for a new piRNA, which shows a characteristic 5’ 10-nt overlap with the original piRNA (ping-pong signature). The new piRNA then reciprocally cleaves complementary RNAs, supplying new piRNA precursors. This piRNA amplification pathway, which depends on the endoribonuclease (slicer) activity of PIWI proteins, is known as the ping-pong cycle^3,5^. In addition, piRNA-guided RNA slicing also recruits an endonuclease named Zucchini (Zuc) downstream of the PIWI protein at the cleavage site. The 3’ fragment of the Zuc-mediated cleavage is then loaded into a new PIWI protein, defining the 5’ end of a new piRNA and recruiting another Zuc to the downstream region. This process, known as the phased piRNA pathway, expands the sequence repertoire of piRNAs beyond the ping-pong sites^9–11^. In both the ping-pong and phased pathways, the 3′ ends of piRNAs are matured by 3′–5′ exonucleolytic trimming by Trimmer in silkworms (PNLDC1 in mammals) and subsequent 2′-*O*-methylation by the methyltransferase Hen1, which stabilizes mature piRNAs^12–14^.

The phased piRNA pathway can be also initiated by specific RNA-binding proteins, for example, Fs(1)Yb in *Drosophila*. A number of protein-coding genes in flies contain Fs(1)Yb-binding sequences in their 3’ untranslated regions (UTR) and generate specific genic piRNAs^15–19^. Accordingly, it is thought that, for transcripts to be processed into piRNAs, they must meet one of the following requirements: integration of the sequence into a piRNA cluster, recognition and cleavage by a pre-exsiting piRNA, or the presence of a specific sequence motif recognized by an RNA-binding protein such as Fs(1)Yb. However, this strict reliance on pre-existing sequence information presents a fundamental chicken-and-egg paradox in germline defense: how does the piRNA system first recognize and target newly invading transposons? Before an invading element is captured by a piRNA cluster to establish “genomic memory,” and before sequence-specific ping-pong amplification can be initiated, the germline must possess a primary mechanism to detect these foreign RNAs without prior sequence knowledge.

In this context, observations in BmN4 cells, derived from silkworm ovaries, offer an intriguing clue. These cells, derived from silkworm ovaries, have both the ping-pong and phased piRNA pathways, making them a valuable model for piRNA research^13,20^. Like silkworm ovaries, BmN4 cells express two PIWI proteins, Siwi and BmAgo3, which participate in the ping-pong amplification of piRNAs^20^. Siwi preferentially associates with piRNAs with uracil at the first position (referred to as 1U bias), whereas BmAgo3-bound piRNAs show a bias toward adenine at the tenth position (referred to as 10A bias)^21,22^. The ping-pong process is thought to occur in a non-membranous liquid-like structure called nuage, which is located around the nucleus^3,23^. Interestingly, we have previously observed that a slicer-deficient Siwi mutant is mislocalized from nuage to P-bodies, another non-membranous structure that contains mRNA decay factors, accompanied by the overproduction of piRNAs derived from genic mRNAs^24^. In fact, these gene-derived piRNAs are produced even under the normal condition, albeit at low levels^24^. However, their biogenesis mechanism remains unclear, because they do not meet the requirements for the known piRNA biogenesis pathways; they are generally uni-stranded in the sense direction, they lack the ping-pong signature, and their sequences do not appear to contain specific motifs for RNA-binding proteins or to be integrated into piRNA clusters^24^. Accordingly, it could be contended that such transcriptome-wide, low-abundance, sense-biased piRNAs are nothing more than mapping or decay artifacts. Alternatively, we hypothesized that these atypical piRNAs might represent the elusive entry point for *de novo* non-self recognition.

Here, we demonstrate that they are bona fide piRNAs that resolve the chicken-and-egg paradox: across silkworms, flies and mice, low levels of sense-strand piRNAs are broadly generated from the transcriptome in proportion to transcript abundance and independently of canonical piRNA biogenesis pathways. In cultured silkworm cells, this pathway generates antiviral piRNAs from *Bombyx mori* latent virus, with piRNA levels directly reflecting viral RNA abundance while efficient ping-pong amplification has not been established. Reanalyses of recently endogenized retroviruses reveal the same entry route in vivo: in mice, AKV-derived piRNAs track viral transcript levels, whereas in koalas, KoRV-A-derived piRNAs have already expanded into ping-pong amplification to some extent. We propose that this inefficient, non-specific, abundance-coupled pathway acts as a naïve surveillance layer in the germline: by sampling transcripts that are abundant and potentially foreign, it initiates a seed pool of piRNAs that can be funneled into more efficient and specific pathways such as ping-pong amplification.

## Results

### Widespread, low-efficiency sense piRNA production across the transcriptome in BmN4 cells

To understand the characteristics of gene-derived sense piRNAs we previously noted^24^, we first mapped piRNA-sized small RNAs (26–32 nt) from BmN4 cells to all annotated genes in silkworms^25^, which include “transposon-like” genes that are homologous to 1,811 previously well-annotated transposons^26^ at the amino acid level. We then performed a metagene analysis to determine the distribution of piRNAs across different gene regions: 5’ UTRs, coding sequences (CDSs), and 3’ UTRs. Many piRNAs were mapped to the antisense strand of genes (Figure 1A, top), which can be attributed to the antisense transcription from transposon-like genes or repetitive sequences in the silkworm genome, such as those in piRNA clusters. We therefore decided to focus on the genes that only have piRNAs mapped to the sense strand. We observed that sense piRNAs are generally produced from all the gene regions, with an apparent peak in the 5’ UTR (Figure 1A, middle). However, this peak was eliminated when we excluded multi-mappable piRNAs (Figure 1A, bottom), suggesting that the peak contained piRNAs homologous to repetitive sequences. In other words, uniquely mapped sense piRNAs showed an almost uniform distribution across 5’ UTRs, CDSs, and 3’ UTRs (Figure 1A, bottom), suggesting that any region of mRNAs can produce gene-derived sense piRNAs. We then performed similar analyses for piRNAs in flies and mice using previously reported PIWI-interacting small RNA datasets^9,10^. Again, after excluding genes with antisense piRNAs and focusing on uniquely mapped piRNAs, we observed largely uniform distributions of gene-derived sense piRNAs across 5’ UTRs, CDSs, and 3’ UTRs (Figure S1A–E), supporting the idea that the whole transcriptome can potentially generate low levels of sense piRNAs. Because UTRs vary widely in length and regulatory content, we focused subsequent analyses on CDS-derived sense piRNAs.

**Figure 1.**
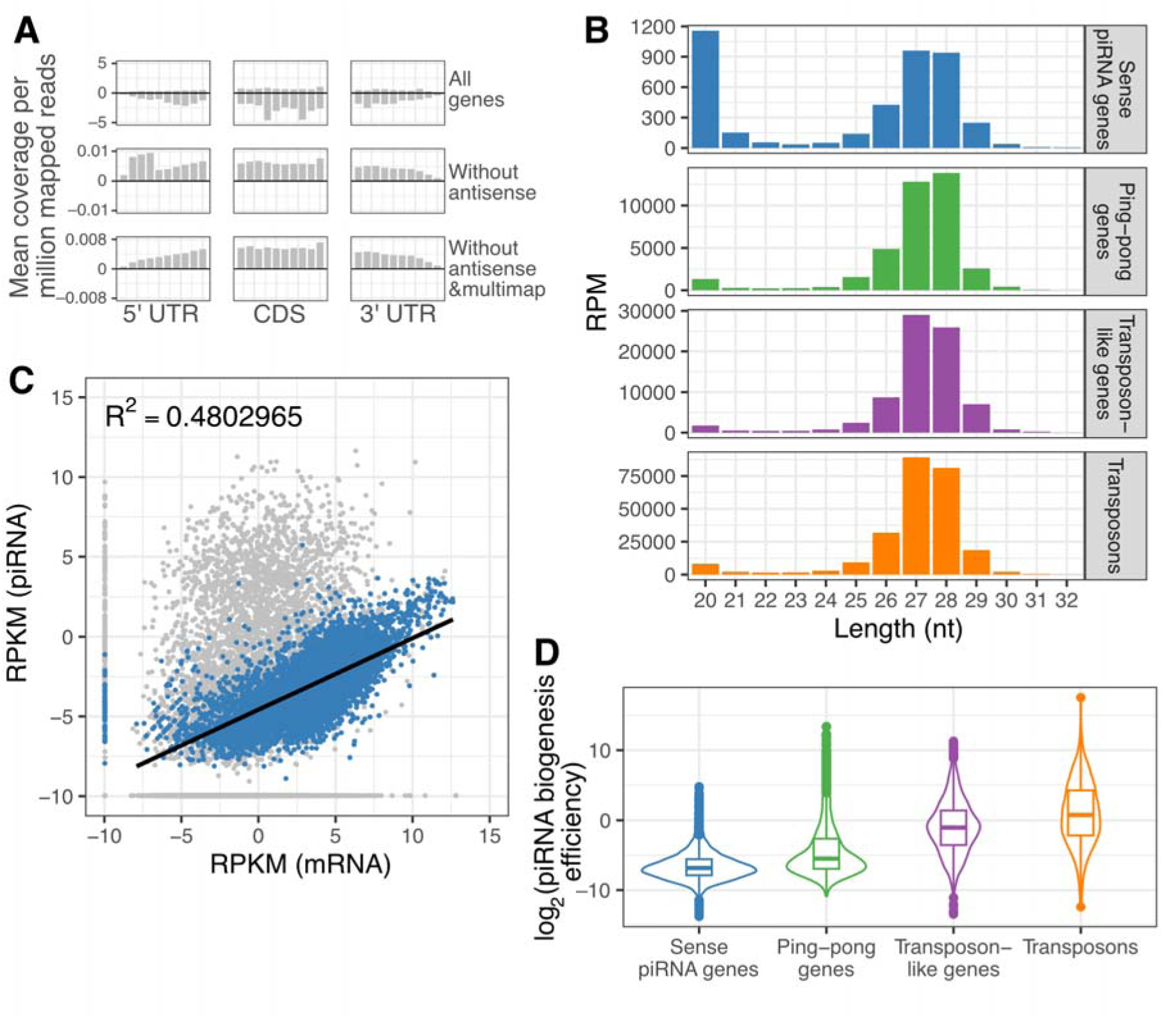
Sense piRNAs are produced from many genes with low efficiency. (**A**) Average piRNA coverage across 5′ UTR, CDS, and 3′ UTR (10 bins each) shown as sense (positive) and antisense (negative); 10,771 of 16,880 genes lacked antisense-mapped piRNAs. (**B**) Length distributions of small RNAs mapped to sense piRNA genes, ping-pong genes, transposon-like genes, and transposons (piRNAs peak at 27–28 nt; siRNAs at 20 nt). (**C**) CDS mRNA versus sense piRNAs (26–32 nt) on log2 axes with a 0.001 pseudocount; sense piRNA genes are highlighted, others are gray; regression fit uses points with log2(mRNA)>0 and log2(piRNA)>0 (adjusted R² shown; see Figure S1H–J). (**D**) piRNA production efficiency [RPKM(piRNA)/RPKM(mRNA)] by category (Center line, median; box, IQR; whiskers, 1.5×IQR; points, outliers).

For the in-depth analysis, we re-sorted the silkworm genes that produce piRNAs from their CDSs (including antisense piRNAs and multi-mappable piRNAs) into three categories: (i) 3,220 genes homologous to the 1,811 well-annotated transposons at the amino acid level (“transposon-like genes”), (ii) 2,033 genes that are not homologous to the transposons but produce sense and antisense piRNAs from the CDSs (“ping-pong piRNA genes”), and (iii) 7,422 genes that are not homologous to the transposons and produce only sense piRNAs from the CDSs (“sense piRNA genes”). We did not detect CDS-derived piRNAs in the remaining 4,205 genes (Figure S1F, Genes without piRNAs). As a reference, we also added piRNAs derived from the CDSs of well-annotated 1,811 transposons^26^ in our analysis (“transposons”).

First, we mapped small RNAs from BmN4 cells to these three gene categories and transposons, and examined their length distribution. We confirmed that the piRNAs derived from the sense piRNA genes have a peak at 27–28 nt, consistent with the canonical piRNA length observed with transposon-like genes, ping-pong piRNA genes, and transposons (Figure 1B). The genes that produce sense piRNAs occupied as much as ∼44% of the entire gene set (Figure S1F). However, these gene-derived sense piRNAs represented only ∼0.7% of the total piRNA pool (Figure S1G), suggesting that their production efficiency is much lower than typical piRNAs generated from transposons, transposon-like genes, and ping-pong piRNA genes. Together, these observations indicate that production of sense piRNAs from ordinary genes is widespread but intrinsically inefficient.

Next, we comprehensively compared the levels of CDS-derived piRNAs and their corresponding source mRNAs across each gene category. We found that transposons, transposon-like genes, and ping-pong piRNA genes produce piRNAs at disproportionally high levels compared to the abundance of their source mRNAs (Figure S1H–J). In contrast, the amount of piRNAs derived from the sense piRNA genes showed a strong proportional correlation to the abundance of their source mRNAs (Figure 1C). These results suggest that many genes are capable of producing sense piRNAs, depending on their mRNA abundance. However, this occurs with a much lower efficiency compared to typical piRNAs (Figure 1D).

### Gene-derived sense piRNAs are bona fide but generated largely independently of canonical pathways

To investigate the characteristics of these gene-derived sense piRNAs, we re-analyzed various silkworm small RNA libraries previously reported. In general, mature piRNAs are 2’-*O*-methyl modified at the 3’ ends by the methyltransferase Hen1, making them resistant to NaIO_4_-mediated oxidation^27^. Indeed, the levels of piRNAs produced from transposons, transposon-like genes, and ping-pong piRNA genes were unaffected by the NaIO_4_ treatment (Figure 2A, top). Similarly, gene-derived sense piRNAs were also resistant to NaIO_4_, indicating that they are also 2’-*O*-methylated (Figure 2A, top).

**Figure 2.**
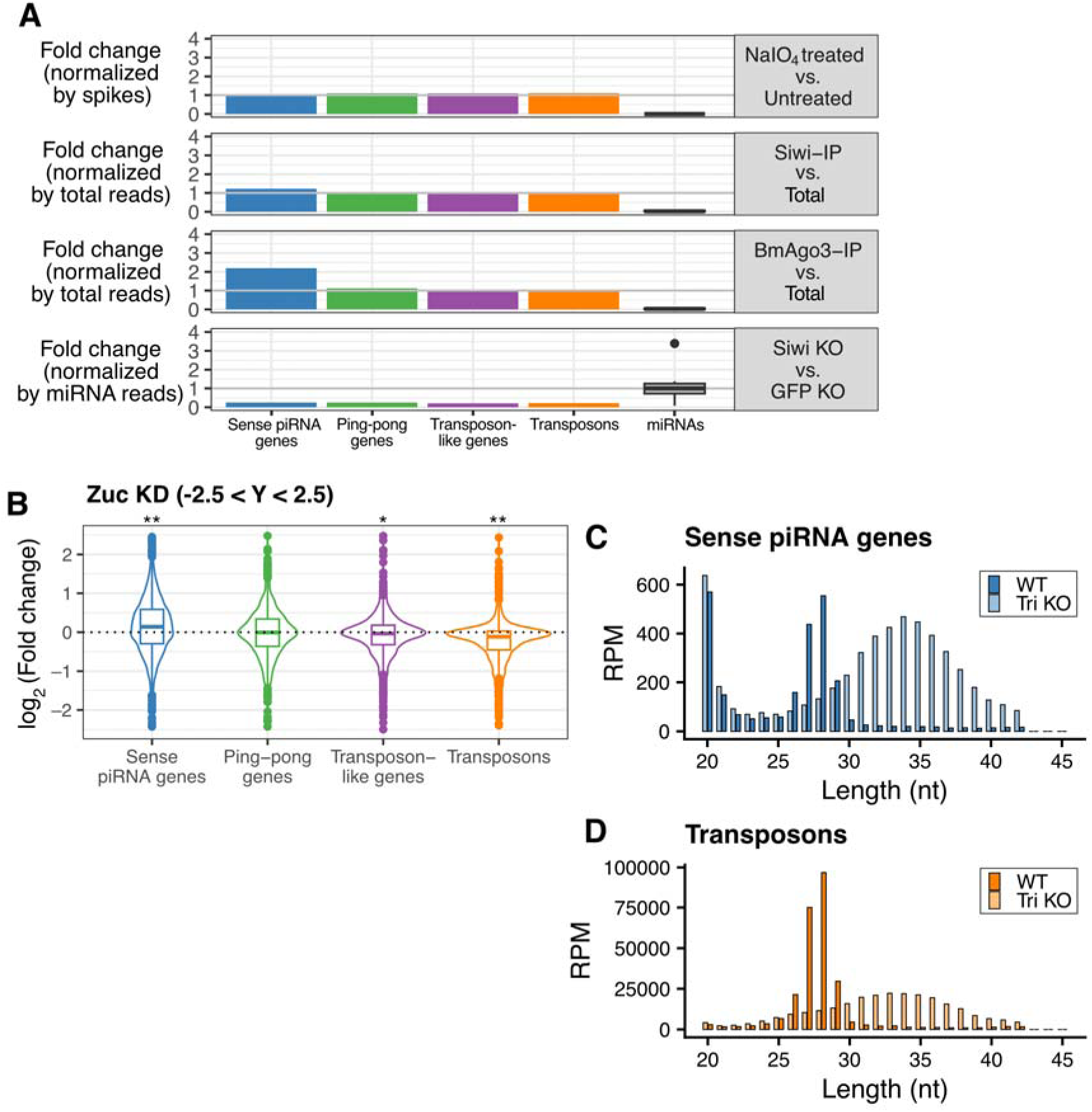
Gene-derived sense piRNAs are bona fide piRNAs. **(A)** Changes in the abundance of gene-derived sense piRNAs, other canonical piRNAs, and miRNAs under biochemical and genetic perturbations. From top to bottom: NaIO treated vs untreated (2′-O-methylation), Siwi-IP vs total library (Siwi binding), BmAgo3-IP vs total library (BmAgo3 binding), and Siwi KO vs control (dependence on Siwi). Totals are shown for piRNAs; miRNAs are shown as boxplots (center line, median; box, IQR; whiskers, 1.5×IQR; points, outliers). **(B)** Violin plots of changes in piRNA abundance upon Zuc knockdown, shown as RPKM(Zuc KD)/RPKM(control KD) for each category; transcripts above the abundance cutoff are included (see Methods). The inset shows values between −2.5 and 2.5. Statistical significance was assessed by one-sample t-test against 0 with Bonferroni correction (four tests). * *P* = 1.5e-6, ** *P* < 8.8e-16. **(C, D)** Length distributions of gene-derived sense piRNAs (**C**) and transposon-derived piRNAs (**D**) in BmN4 cells comparing wild type and Trimmer knockout; reads are normalized by total genome-mapped reads.

To confirm that these small RNAs are bona fide piRNAs, we examined their association with PIWI proteins. piRNAs deriving from transposons, transposon-like genes, and ping-pong piRNA genes were significantly enriched by both Siwi– and BmAgo3-immunoprecipitation (IP), compared to microRNAs (miRNAs) that are known to interact with AGO-subfamily proteins (Figure 2A, middle). Interestingly, gene-derived sense piRNAs were even more enriched by Siwi– and BmAgo3-IP, presumably because they are too rare to be quantitatively detected without being concentrated by IP. On the other hand, Siwi– and BmAgo3-bound gene-derived piRNAs showed a strong correlation with the abundance of their source mRNAs, suggesting that they are generally associated with PIWI proteins without specific selection (Figure S2A, B). Conversely, gene-derived sense piRNAs were significantly depleted by the transient knockout (KO) of Siwi, like typical piRNAs (Figure 2A, bottom). We concluded that small RNAs produced from sense piRNA genes are bona fide piRNAs, bearing 2’-*O*-methyl modification at the 3’ end and interacting with PIWI proteins.

Given that these sense piRNAs from genes lack antisense ping-pong pairs, a plausible explanation is that they are produced by the Zuc-mediated phased piRNA biogenesis pathway. To examine the requirement of Zuc in the production of gene-derived sense piRNAs, we investigated the change in their expression upon the knockdown (KD) of Zuc in BmN4 cells (Figure 2B). Transposon-derived piRNAs were significantly reduced by Zuc KD, suggesting that they are at least in part generated by the phased pathway in addition to the ping-pong pathway. piRNAs derived from transposon-like genes and ping-pong piRNA genes were less affected by the Zuc KD, suggesting that they are mainly generated by the ping-pong pathway. In contrast, gene-derived sense piRNAs were rather upregulated by Zuc KD, suggesting that their production does not require either the ping-pong cycle or Zuc-dependent phased processing, at least in silkworm BmN4 cells (see Discussion). Given that Trimmer matures piRNAs by 3′–5′ exonucleolytic trimming whether their precursor 3′ ends are generated by Zuc-dependent cleavage or by ping–pong–associated slicing^13^, it is likely capable of processing 3′-extended ends generated by any nucleolytic events. Such robust Trimmer activity could therefore account for the apparent dispensability of Zuc in BmN4 cells. Indeed, Trimmer knockout caused a broad 3′ extension of piRNAs from both gene-derived sense transcripts and transposons in BmN4 cells (Figure 2C, D). Therefore, 3′–5′ exonucleolytic trimming by Trimmer specifies the mature 3′ ends of gene-derived sense piRNAs, much as it does for canonical transposon-derived piRNAs.

### Gene-derived sense piRNAs are also produced in flies and mice

To investigate whether gene-derived sense piRNAs are specific to silkworms or a more general phenomenon, we performed similar analyses using publicly available datasets and CDSs from *Drosophila* and mice. We found that gene-derived sense piRNAs are also generated with low efficiency in *Drosophila*; they are associated with all three fly PIWI proteins (Aubergine [Aub], Ago3, and Piwi) and their abundance is proportional to their source mRNA levels (Figure 3A–C). As observed in silkworms, transposons and some ping-pong piRNA genes in flies also produced disproportionally abundant piRNAs compared to their source mRNAs (Figure S3A–F; note that the gene model of *Drosophila* is so well-annotated that transposon-like genes are virtually excluded). Moreover, gene-derived sense piRNAs associated with Piwi and Aub showed typical length distributions peaking at ∼25–26 nt, although the length peak of those associated with Ago3 was less clear (Figure 3D). Similarly in mice, we found that gene-derived sense piRNAs are associated with MILI or MIWI2 with typical length peaks at ∼26 nt and ∼28 nt, respectively (Figure 3E). Their abundance is proportional to their source mRNAs (Figure 3F, G) and lower than piRNAs deriving from ping-pong piRNA genes and transposons (Figure S3G–J). These data support the idea that genes generally produce low levels of piRNAs in flies and mice, like in silkworms.

**Figure 3.**
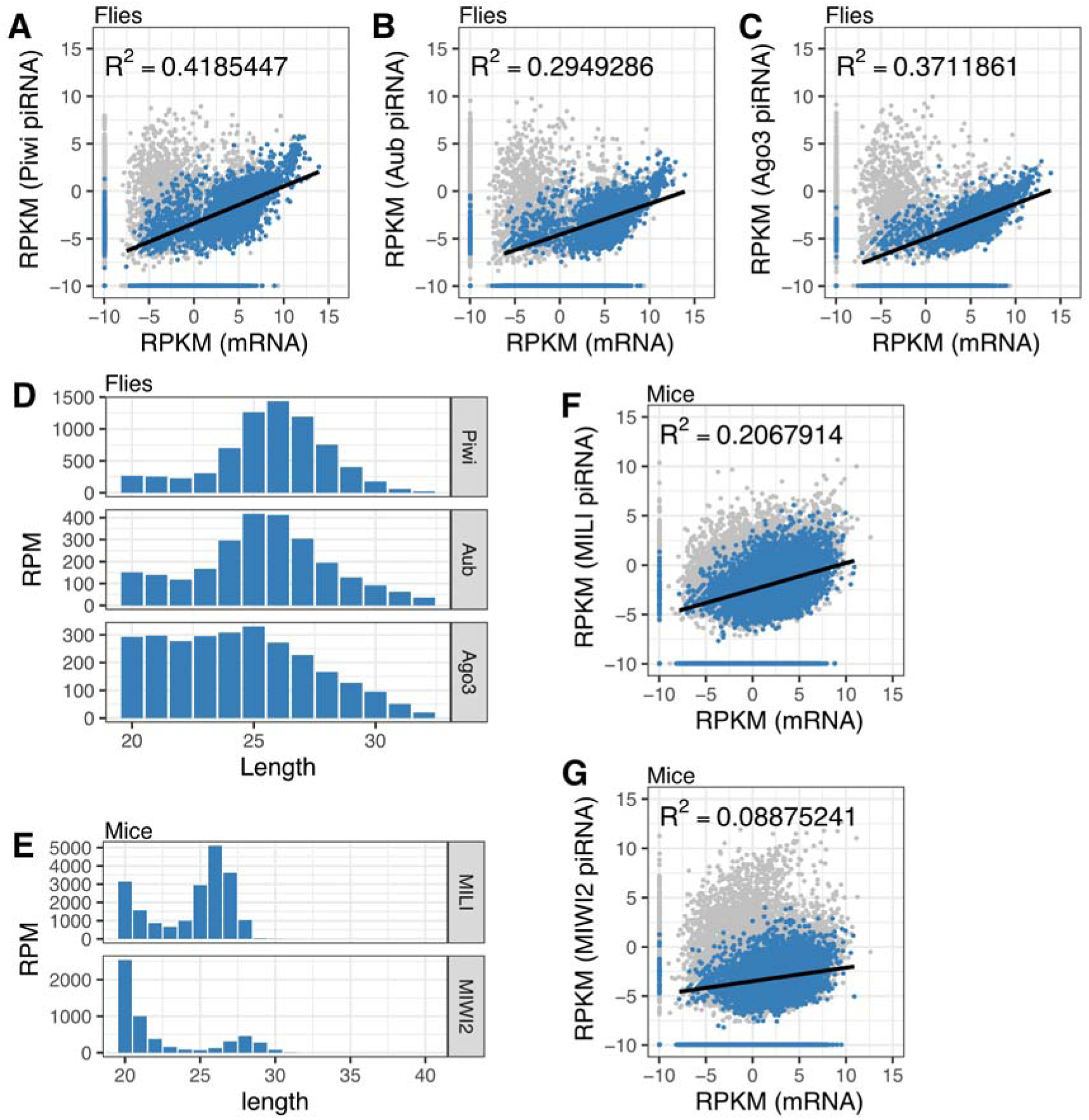
Gene-derived piRNAs in flies and mice. **(A–C)** Scatter plots of ovarian mRNA abundance and sense-mapped piRNAs bound to Piwi (**A**), Aub (**B**), and Ago3 (**C**) in *Drosophila melanogaster*. Genes with no antisense-mapped piRNAs in all three IP libraries (n = 7,106) are plotted. Axes are log2 with a 0.001 pseudocount; regression lines are fit using points with log2(mRNA) > 0 and log2(piRNA) > 0, and adjusted R² is shown. **(D)** Length distributions of gene-derived piRNAs bound to Piwi, Aub, and Ago3. The peak lengths (Piwi, 26 nt; Aub, 25 nt; Ago3, 25 nt) match the canonical size profiles previously reported for transposon-derived piRNAs bound to these PIWI proteins^10^. **(E)** Length distributions of gene-derived piRNAs bound to MILI and MIWI2 in mouse primary spermatocytes. The peak lengths (MILI, 26 nt; MIWI2, 28 nt) are consistent with the reported size profiles of canonical (transposon-derived) piRNAs for these proteins^9^. **(F,G)** Scatter plots of primary spermatocyte mRNA abundance and MIWI2– (**F**) or MILI-bound (**G**) sense piRNAs mapped to genes. Genes lacking antisense piRNA mapping (n = 20,656) are plotted; axes, pseudocount, and regression criteria are as in **(A–C)**.

Long non-coding RNAs (lncRNAs) are particularly well annotated in the mouse RefSeq genome database^28^. Therefore, we investigated if sense piRNAs are also produced from mouse lncRNAs. We extracted 2,056 lncRNAs that produce only sense piRNAs (sense piRNA lncRNAs) and 1,392 lncRNAs that produce antisense piRNAs (ping-pong piRNA lncRNAs), and then mapped piRNAs to them. We found that, as observed with gene-derived piRNAs, lncRNA-derived sense piRNAs are produced with comparable efficiency to gene-derived sense piRNAs, and their abundance is proportional to their sources (Figure S3K–N). Taken together, these results indicate that low-level sense piRNA production occurs across the transcriptome, rather than being restricted to protein-coding genes.

### Abundant virus-derived piRNAs are generated by the same pathway as gene-derived piRNAs in silkworm cells

A positive-sense single-stranded RNA virus called *Bombyx mor*i latent virus (BmLV) is known to persistently infect BmN4 cells, where its replication is suppressed by piRNAs^29–31^. BmLV-derived piRNAs primarily originate from the extremely abundant sense-stranded sub-genomic RNAs and target the transiently synthesized antisense strand during viral replication^30^. However, it remains unknown how BmLV-associated piRNAs are produced. To answer this question, we first quantified BmLV-derived piRNAs and their source mRNAs (RNA-dependent RNA polymerase [Rdrp), Coat protein [Cp), and P15), as we did for gene-derived sense piRNAs (Figure 4A). We found that both piRNAs and mRNAs from BmLV are extremely abundant but the ratio between them is similar to that of general gene-derived sense piRNAs and their source mRNAs in BmN4 cells (Figure S4A). We have previously reported that overexpression of a Siwi mutant (Siwi-D670A) that disrupts its slicer activity causes selective upregulation of gene-derived piRNAs, presumably because of the suppression of the slicer-dependent ping-pong pathway^24^. By reanalyzing this dataset, we found that BmLV-derived piRNAs are also significantly increased by Siwi-D670A (Figure 4B). In contrast, overexpression of Siwi-D670A did not cause any changes in the expression levels of BmLV-derived small interfering RNAs (siRNAs), which are mainly produced from the genomic region of BmLV and contribute to virus silencing together with BmLV-derived piRNAs^30,31^. Moreover, BmLV-derived piRNAs were resistant to NaIO_4_ treatment, bound to PIWI proteins sensitive to Siwi knockout, and unaffected or slightly upregulated by Zuc knockdown (Figure 4C, S4B, C). These results suggest that BmLV-derived sense piRNAs are produced by the same pathway as gene-derived sense piRNAs; BmLV-derived piRNAs are abundant simply because their source mRNAs are abundant.

**Figure 4.**
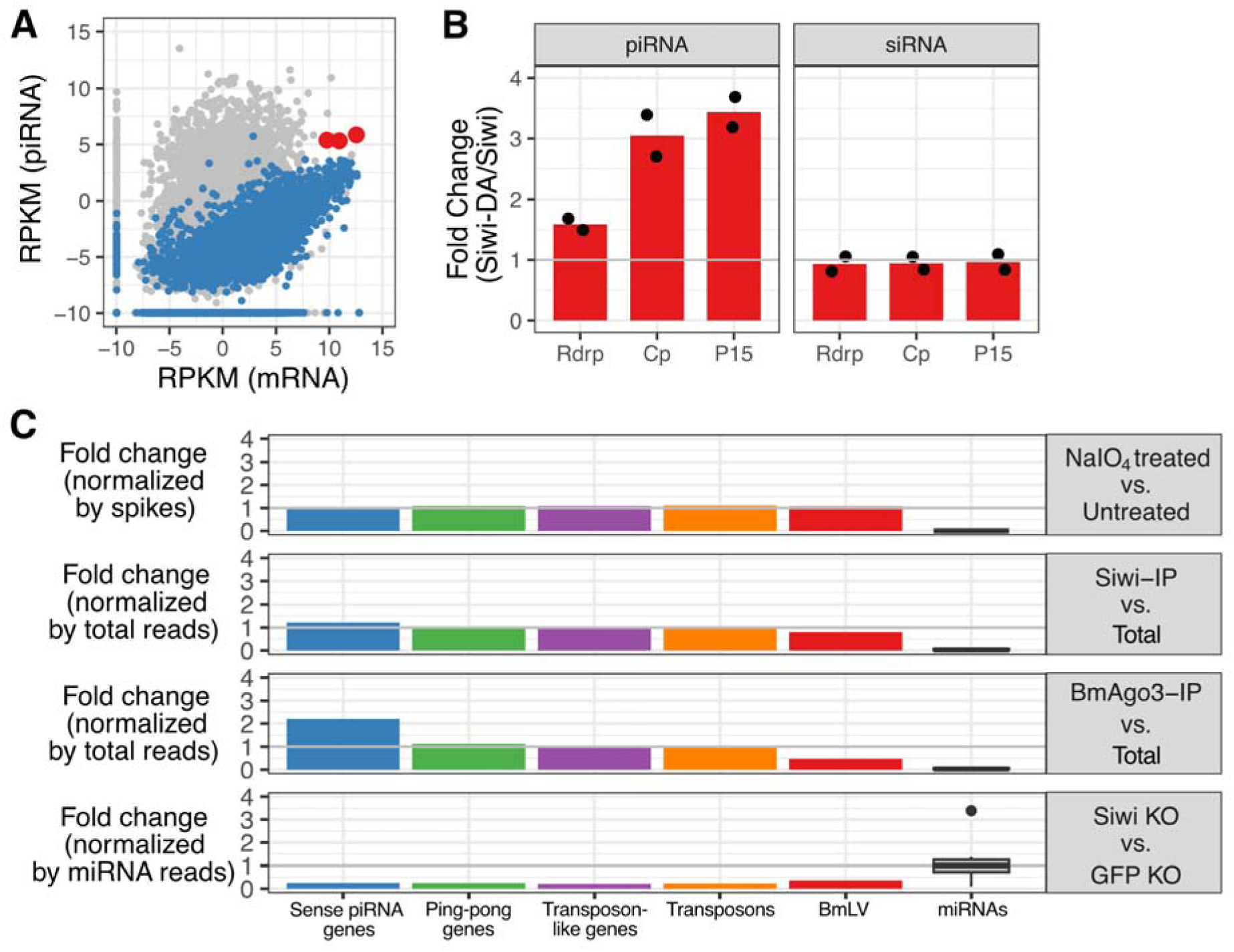
BmLV-derived piRNAs are produced by the same pathway as gene-derived piRNAs. **(A)** Scatter plot of sense piRNA abundance (26–32 nt) plotted against CDS mRNA abundance (log2; 0.001 pseudocount). Sense piRNA genes are shown in blue, the three BmLV ORFs are highlighted in red. **(B)** Expression of catalytically inactive Siwi (Siwi-DA) increases BmLV-derived 26–32-nt piRNAs, whereas 20-nt siRNAs do not; two replicate libraries are shown with means, and a horizontal gray line at y = 1 indicates no change. **(C)** NaIO treated vs untreated, Siwi-IP vs total, Ago3-IP vs total, and Siwi KO vs control assess BmLV piRNAs relative to other canonical piRNAs and miRNAs; plotting conventions are as in Figure 2A.

### In vivo evidence for naïve piRNA biogenesis during retroviral endogenization

To examine whether our proposed model also applies in vivo—especially under conditions where piRNA-mediated silencing is not yet fully established, as in recent retroviral invasions—we reanalyzed small RNA datasets from koala and mouse testes^32,33^. In mice, AKV murine leukemia virus—a recently endogenized retrovirus in the AKR strain—produces sense piRNAs without apparent ping-pong signatures (Figure S5A)^32,34^. Moreover, our reanalysis showed that the piRNA/mRNA ratios of AKV-derived piRNAs remain close to the median level for those derived from sense piRNA genes (Figure 5A). We concluded that this represents a situation similar to that observed for BmLV in BmN4 cells, in which abundance-coupled, sense piRNA biogenesis is at work (Figure 4).

**Figure 5.**
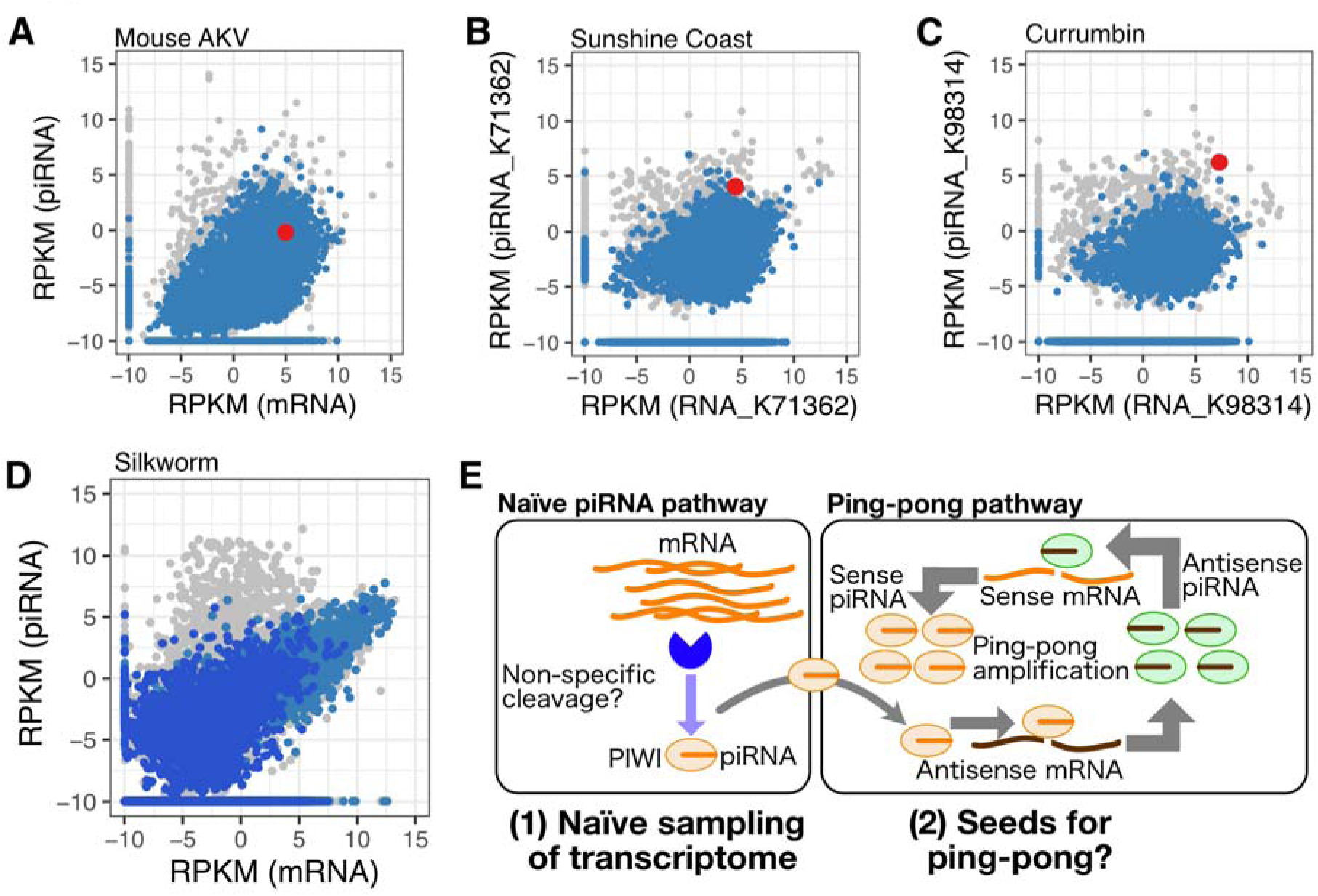
In vivo evidence for naïve piRNA biogenesis during retroviral endogenization. **(A–C)** Scatter plots show the relationship between mRNA abundance and piRNAs (≥26 nt) in mouse testes (**A**), koala testes from Sunshine Coast (**B**), and koala testes from Currumbin (**C**). In each sample, mRNAs and piRNAs were mapped to genes in the sense orientation. Genes with no antisense piRNA mapping are colored **blue** (mouse, n = 19,551; koala, n = 43,042). AKV is highlighted in **red** in **(A)**. KoRV-A is highlighted in **red** in **(B)** and **(C).** Axes are log2 with a 0.001 pseudocount added to both axes. **(D)** Exon–intron comparison of piRNA production in BmN4 cells: for 2,080 genes with exclusively sense piRNA mapping across the gene body (including introns), mRNA RPKM (x-axis) versus sense piRNA RPKM (y-axis) is plotted for concatenated exonic (**blue**) and intronic (**navy**) regions (definitions in Methods). **(E)** Model: abundant transcripts are fragmented by putative nonspecific nuclease(s) and loaded into PIWI proteins as naïve piRNAs with low efficiency proportional to mRNA abundance; virus-derived piRNAs can target and repress viral antisense mRNAs and may seed later amplification upon invasion of new elements.

In contrast, koala retrovirus A (KoRV-A)—a recently acquired gammaretrovirus still spreading through vertical and horizontal transmission^32,33,35,36^—shows evidence of progression into ping-pong amplification. Yu et al.^33^ recently reported that koala populations south of the Brisbane River (e.g., Currumbin) are characterized by sense-biased piRNA production without efficient viral silencing, whereas a subpopulation north of the river (e.g., Sunshine Coast) exhibits strongly suppressed KoRV-A expression and more balanced sense/antisense piRNAs. However, our reanalysis detected clear 10-nt ping-pong signatures of KoRV-A-derived piRNAs not only in Sunshine Coast animals but also in Currumbin individuals, and their piRNA/mRNA ratios often exceeded those of typical sense piRNA genes (Figure 5B, C and Figure S5B–F). Consistent with this, an earlier study by Yu et al.^32^ had already detected a significant ping-pong signature in at least one Currumbin individual (Figure S5B in^32^). Together, these observations indicate that KoRV-A piRNA responses in these koala populations are no longer confined to an abundance-coupled baseline (as in BmLV or AKV-MLV) but have progressed—albeit to varying extents—into ping-pong amplification.

These findings support a model in which the abundance-coupled sense piRNA pathway acts as a constitutive, transcriptome-wide mechanism that seeds silencing by generating low-level sense piRNAs. Once both sense and antisense transcripts are sufficiently available, this entry route can transition into more efficient suppression through ping-pong amplification and additional epigenetic regulation. We refer to this initial, abundance-coupled state as “naïve piRNA biogenesis”— naïve in the sense that, much like naïve immune cells, it is initially non-selective and low-efficiency but provides the raw material that can be shaped into a highly specific and robust defense. The naïve piRNA biogenesis pathway functions as a latent surveillance system, enabling the germline to detect and initiate silencing of aberrantly abundant or newly emerging genomic elements. We propose that such constitutive low-level piRNA production provides a critical foundation for the adaptability and robustness of the piRNA-mediated genome defense system.

### Naïve piRNA biogenesis is distinct from other small RNA pathways

Yu et al.^33^ reported that “innate” piRNAs for KoRV-A are preferentially generated from unspliced, intron-retaining RNAs. To examine whether this applies more generally to gene-derived naïve sense piRNAs, we quantified intronic piRNA/mRNA ratios in silkworms after excluding genes that contained any antisense-oriented piRNA reads across the gene body, including introns. This filtering yielded 2,080 genes, for which we plotted intronic piRNA/mRNA ratios. Introns largely followed the same trend as exons, albeit at lower absolute abundance (Figure 5D). In other words, although introns yielded fewer piRNAs overall, their piRNA/mRNA relationship was essentially an extension of exon-derived piRNAs. These observations support a model in which naïve piRNA biogenesis samples broadly from expressed transcripts, rather than preferentially targeting intron-retaining or mis-spliced RNAs.

To further distinguish naïve piRNAs from other small RNAs produced under distinct regulatory mechanisms, we revisited a recent report in *Drosophila melanogaster* testes, piRNAs are produced from CDSs, but not from 5’ or 3’ UTRs, in a manner dependent on Ago2, an AGO protein that loads siRNAs and some miRNAs, and Dicer-2, the core biogenesis factor for siRNAs^37^. Our reanalysis confirmed that these CDS-piRNAs are markedly down-regulated by the knockdown of Ago2 or Dicer-2 (Figure S5G, H). In contrast, gene-derived naïve sense piRNAs were only slightly decreased by Ago2 KD and remained unchanged by Dicer-2 KD, like piRNAs derived from ping-pong genes or transposons (Figure S5G, H), suggesting that CDS-piRNAs and naïve piRNAs are distinct entities.

Finally, to assess whether naïve piRNA production requires canonical piRNA factors, we examined two systems lacking complete piRNA machinery. OSC cells, derived from the ovarian somatic cells of *Drosophila*, lack the ping-pong pathway and are thought to produce piRNAs solely via the Fs(1)Yb-dependent “primary” pathway^38^. *Drosophila* embryo-derived Kc167 cells express Piwi and Aub, but they lack the essential components required to produce transposon-derived piRNAs, including factors required for piRNA cluster transcription; accordingly, piRNAs in Kc167 cells are mainly derived from mRNAs and tRNAs^39^. Nevertheless, reanalysis of these datasets revealed low-frequency, sense-biased piRNAs whose abundance scaled with gene expression (Figure S5I–K), indicating that abundance-coupled naïve piRNA biogenesis persists even in the absence of a functional ping-pong pathway or cluster-dependent production.

## Discussion

Our study identifies a conserved, abundance-coupled mode of piRNA production that generates sense-biased piRNAs broadly from the transcriptome in silkworms, flies, and mice (Figure 1, 3). These “naïve” piRNAs bear molecular hallmarks of bona fide piRNAs (Figure 1B, 2A, C) yet operate independently of canonical ping-pong and Zucchini-dependent phasing (Figure 1A, 2B). The same mechanism engages viral RNAs: BmLV in silkworm BmN4 cells and AKV in mice remain near the abundance-coupled baseline (Figure 4A, 5A and Figure S5A), while KoRV-A has already progressed to ping-pong amplification (Figure 5B, C and Figure S5B–F). Rather than replacing canonical pathways, this background activity supplies a low-level pool of piRNA sequences that can be amplified when complementary antisense transcripts and amplification machinery become available. Functionally, this abundance-coupled route behaves like a basal surveillance layer: it passively samples the transcriptome for abundant and potentially foreign RNAs and thereby seeds an initial pool of sequences that can be recruited into more potent silencing programs.

Our recent work has shown that the piRNA sequence repertoire is autonomously optimized by competition among adjacent ping-pong sites, a process that favors sequences that target and silence transposons more effectively^40^. Optimization is thought to occur as new piRNAs emerge randomly and compete with existing ones nearby: more efficient piRNAs are preferentially amplified, while less effective sequences decline and are eventually eliminated. What has been unclear is the source of those novel sequences. We propose that naïve, abundance-coupled production supplies a continuous, low-level stream of candidate piRNAs across the transcriptome. By feeding these new sequences into the competitive dynamics of ping-pong, naïve production continually drives optimization of the repertoire, ensuring that the piRNA pool can keep pace with rapidly mutating transposons and maintain effective silencing.

Notably, our conclusions converge with those of Handler et al. (accompanying manuscript), who independently find that piRNA biogenesis is built on a naïve sampling framework that non-selectively draws on available transcripts and is later refined by context-dependent specificity modules, such as Fs(1)Yb-mediated recognition of uridine-rich precursor RNAs derived from adenine-rich retrotransposon sequences in *Drosophila* ovarian somatic cells (OSC). In our analysis, gene-derived sense piRNAs are rather upregulated upon Zuc KD (Figure 2B and Figure S2C), indicating that—at least in silkworm BmN4 cells—naïve piRNAs do not require the Zuc-dependent phasing pathway for production. By contrast, Handler et al. find a stronger dependence on Zuc and Armitage (Armi) in *Drosophila* OSC for generating naïve piRNAs. It has been shown that, in silkworms, the 3′–5′ exonuclease Trimmer efficiently trims extended piRNA 3′ ends generated downstream of both Zuc-mediated and ping-pong–mediated cleavage^12,13^, and we found that naïve piRNAs are likewise matured by Trimmer (Figure 2C, D). Trimmer (PNLDC1) is broadly conserved in mammals and worms but is absent in *Drosophila*, which instead relies on another 3′–5′ exonuclease, Nibbler, to trim piRNA 3′ ends; nevertheless, Nibbler can trim only a few nucleotides, and many fly piRNAs still require Zuc for 3′-end formation^10,11,41^. This difference in 3′-end processing activities plausibly explains why naïve piRNAs do not require Zuc in silkworm BmN4 cells, whereas *Drosophila* OSC show a stronger Zuc/Armi dependence in the accompanying study.

These observations, together with our broader analyses, narrow the possibilities for how naïve piRNAs are generated. We see no strong enrichment for intron-retaining or mis-spliced RNAs—intronic signals simply mirror the exon-derived piRNA/mRNA relationship at lower abundance (Figure 5D)—and naïve piRNAs are distinct from CDS-restricted piRNAs in *Drosophila* testis, which collapse after Ago2 or Dicer-2 knockdown, whereas naïve species persist (Figure S5G, H). Moreover, low-level, abundance-coupled sense piRNAs are detectable in multiple cellular contexts, including those that lack a functional ping-pong cycle (OSC; Figure S5I) or cluster-dependent piRNA production (Kc167; Figure S5J, K). We therefore infer that a broadly acting source of RNA fragments—largely non-selective at the level of which transcripts are sampled—can feed into different biogenesis modules rather than relying on any single canonical route. In systems such as silkworms, where Trimmer/PNLDC1 is likely able to efficiently trim 3′ extensions of various length generated by diverse upstream nucleolytic events^12,13^, broadly non-specific, stochastic RNA degradation may play a major role in supplying substrates for the naïve pathway. By contrast, in *Drosophila*, which lacks Trimmer/PNLDC1, production of naïve precursors is likely to rely more directly on Zuc-mediated endonucleolytic cleavage to generate suitably sized entry fragments for abundance-coupled sampling of available transcripts. Despite these mechanistic differences, both our data and the accompanying study point to the same overall architecture: a broadly sampling, abundance-coupled naïve layer that is wired into context-specific modules (such as Fs(1)Yb or ping-pong) to generate efficient, sequence-specific genome defense.

A mechanistic clue to how naïve piRNAs are produced comes from our previous finding that overexpression of the slicer-deficient Siwi-D670A mutant increases sense-strand naïve piRNAs from both genes^24^ and BmLV (Figure 4B). Siwi-D670A mislocalizes to P-bodies, forming more solid-like assemblies than the normal, liquid-like nuage. Although P-bodies are generally viewed as mRNA storage/decay sites^42,43^, they also contain a subset of piRNA biogenesis factors in germ cells^24,44,45^. We hypothesize that, under normal conditions, Siwi dynamically shuttles between nuage and P-bodies, whereas Siwi-D670A loses this mobility and becomes immobilized in P-bodies, thereby channeling P-body–stored mRNAs into the piRNA pathway. In this scenario, the non-specific RNA fragmentation that generates naïve piRNAs may occur within P-bodies even under normal conditions. Consistent with canonical maturation, naïve piRNAs are trimmed by Trimmer (Figure 2C, D) and 2′-*O*-methylated at their 3′ ends (Figure 2A). Thus, once fragments are loaded onto a PIWI protein, they appear to complete maturation on the mitochondrial outer membrane via the standard processing pathway. Conversely, piRNA-guided recognition of complementary targets and initiation of ping-pong cleavage are generally thought to occur in perinuclear nuage^3,5,23,46^. Viewed this way, dynamic subcellular compartmentalization among P-bodies, nuage, and mitochondrial surface may act as a fidelity gate that buffers the naïve surveillance layer from the high-gain ping-pong amplifier, minimizing crosstalk and ensuring robust amplification is engaged only when complementary, non-self signals are present.

## Materials and Methods

### Cell culture

BmN4 cells were cultured at 27°C in IPL-41 medium (Applichem) supplemented with 10% fetal bovine serum.

### Sequence analysis of small RNAs

Small RNA sequences from BmN4 cells (DRR181866) were reported previously^13^. For informatics analysis, the 3′-adaptor sequences were identified and removed by cutadapt 2.6 with Python 3.7.0 with “--minimum-length 20” parameter. Reads that could be mapped by bowtie 1.2.3^47^ to the *Bombyx mori* genome^25^ or *Drosophila melanogaster* genome r6.27 from FlyBase^48^ or mouse genome GRCm38.p5 from NCBI, allowing up to two mismatches, were used to calculate the mapping rate and to normalize each library. Mapping to the BmLV genome (GenBank *AB624361*.1) was performed under a similar condition but the reads were not used for normalization. In the comparison between the silkworm NaIO_4_-treated library and its control, small RNA reads were normalized by the spike sequences (Supplementary Table 1) introduced into the library. In the comparison of the Siwi KO library with its control, normalization was performed using the median read of miRNAs (Supplementary Table 1) present in the library, as in the previous analyses^12,49^. Of the 96,858 mouse CDSs, the top-listed isoform in the FASTA file for each gene was treated as a representative, and 22,858 CDSs were used for analysis. Reads that could be mapped by bowtie 1.2.3^47^ to the *Bombyx mori* gene model^25^ or *Drosophila melanogaster* CDSs dataset r6.27 from FlyBase^48^ or mouse CDSs dataset GRCm38.p5 from NCBI, allowing up to two mismatches were used for downstream analyses. SAM files were converted to BAM files by SAMtools^50^, then to BED files, and the mapped reads of each gene and transposons were calculated by BEDTools^51^. Those reads with a length between 26 nt and 32 nt were defined as piRNAs.

Silkworm “sense piRNA genes” were defined as genes with ≥1 sense-mapped piRNA read (26–32 nt) and 0 antisense-mapped piRNA reads in the corresponding dataset. Silkworm ping-pong genes were defined as genes with >1 antisense-mapped piRNA read in the total small RNA library of the GFP knockout BmN4 cells. Silkworm transposon-like genes (i.e., genes containing transposon-related sequences) were defined as in the previous paper^24^. Specifically, TBLASTX was performed using 1,811 transposon sequences (26) as the database and silkworm gene model sequences as the queries, and genes having an E-value of 1e-50 or less were defined as transposon-like genes. Transposon-derived reads were quantified by mapping reads to the 1,811 transposon sequences^26^ under the same bowtie settings (≤2 mismatches) and counting mapped reads per transposon.

For the Drosophila and mouse data, we used only the top-listed splice variants from the CDSs FASTA files. In addition, we excluded those genes whose piRNAs mapped to the antisense strands with more than 1 read in the three libraries, Piwi-IP, Aub-IP, and Ago3-IP for Drosophila and in the two libraries, MILI-IP and MIWI-IP for mice.

For analyses comparing positional piRNA coverage across transcript features, 5′ UTRs, CDSs, and 3′ UTRs were each divided into 10 equal-length bins per gene. Sense and antisense coverages were computed separately and plotted with sense values on the positive axis and antisense values on the negative axis.

### Sequence analysis of mRNAs

mRNA reads that could be mapped by Hisat 2.1.0^47^ to the *Bombyx mori* genome^25^ or *Drosophila melanogaster* genome r6.27 from FlyBase^48^ or mouse genome GRCm38.p5 from NCBI with –k 3 option were used to calculate the mapping rate and to normalize each library. Mapping to the BmLV genome (GenBank *AB624361*.1) was performed under a similar condition but the reads were not used for normalization. SAM files were converted to BAM files by SAMtools^50^, then to BED files, and the mapped reads of each gene and transposons were calculated by BEDTools^51^. or exon–intron comparisons, exonic and intronic segments were concatenated separately for each gene. mRNA and piRNA RPKMs were calculated for the concatenated exonic and intronic regions independently. Transposon-derived mRNA reads were quantified by mapping reads to the 1,811 transposon sequences^26^ under the same settings and counting mapped reads per transposon. Results were imported into R and graphed.

### Analysis of small RNA lengths

Reads mapped to genes, transposons, and BmLV were extracted from FASTA files and calculated the length distributions of the mapped piRNAs. The alphabetFrequency function of the biostrings package in R was used for the calculation. When comparing libraries, length distributions were normalized by total genome-mapped reads.

### Knockdown of Zucchini, RNA extraction, and small RNA-seq

Preparation of a small RNA library from Zuc KD cells was performed as described previously^12^. For double-stranded RNA (dsRNA) preparation for KD, template DNAs were prepared by PCR using primers containing T7 promoter listed in Supplementary Table 1. dsRNAs were transcribed using T7 Scribe Standard RNA IVT Kit (Cell Script) and purified with MEGAclear Transcription Clean-Up Kit (Thermo Fisher/Invitrogen). For dsRNA transfection, 5 μg of dsRNAs were transfected into BmN4 cells (6 × 10^5^ cells per 10 cm dish) with X-tremeGENE HP DNA Transfection Reagent (Sigma). dsRNAs were repeatedly transfected every 3 days for four times. Total RNAs were prepared by TRI Reagent (Molecular Research Center, Inc.). Small RNA libraries were prepared from ∼20–45 nt total RNAs using Small RNA Library Preparation kit (Illumina) and analyzed by Illumina HiSeq 2500 platform.

### Data and code availability

The sequencing data obtained in this study are available under the accession number DRA017491 (Zuc KD). Small RNA sequences from control BmN4 cells (DRR241076), Siwi-knockout cells (DRR241078), wild type and Trimmer-knockout cells (DRR181866 and DRR181867) untreated (DRR181866), NaIO4 treated (DRR181868), Siwi-IP and BmAgo3-IP piRNA sequences (DRA017492), mRNA sequence from control BmN4 cells (DRR097455) were reported previously^12,13,24,30,52^. Drosophila small RNA sequences of Piwi-IP (SRR1746863), Aub-IP (SRR1746864) and Ago3-IP (SRR1746865), Drosophila mRNA sequences (SRR2147092) were reported previously^10,53^. Mouse unoxidized small RNA sequences of MILI-IP (SRR5304361), MIWI2-IP (SRR5304364) from P0 testis, mouse mRNA sequences (SRR7760359, SRR7760371 and SRR7760372) from P0 testis were reported previously^54,55^. mRNA-seq derived from the OSC of Drosophila (SRR9158291) and small RNA-seq of Piwi-IP (SRR9158321), mRNA-seq from Kc167 cells (SRR030164) and small RNA-seq from Piwi-IP and Aub-IP (SRR4429377, SRR4429376) have been previously reported^39,56,57^. mRNA-seq from Sunshine Coast koala testes (SRR30589450 and SRR30589454) and from Currumbin koala testes (SRR8708139, SRR8708140 and SRR30589446), as well as oxidized small RNA-seq from Sunshine Coast (SRR30589597 and SRR30589600) and Currumbin testes (SRR8708150, SRR8708151 and SRR30589589), have been previously reported^32,33^. mRNA-seq (SRR9080662) and small RNA-seq (SRR9080667, SRR9080668, SRR9080669) derived from mouse testes have been previously reported^32^. All code required for bioinformatics analysis in this paper is available at https://github.com/keishoji/mRNApiRNA.

## Acknowledgments

We thank all the members of the Tomari laboratory for discussion and critical comments on the manuscript. We thank Dr. Natsuko Izumi of The University of Tokyo for preparation of the Zuc KD library and careful and critical comments on this paper. We thank Dr. Masashi Iwanaga of Utsunomiya University and Dr. Susumu Katsuma of The University of Tokyo for their fruitful discussion. We disclose that AI-assisted tools were used to improve the clarity and readability of the manuscript. This work was supported in part by Grant-in-Aid for Scientific Research (S) (18H05271 to Y.T.), Grant-in-Aid for Scientific Research (A) (23H00364 to Y.T.), Grant-in-Aid for JSPS Fellows (17J02408 to K.S.), Grant-in-Aid for Transformative Research Areas (A) (24H02278 to K.S.), Grant-in-Aid for Scientific Research (B) (24K01769 to K.S.).

## Author contributions

Conceptualization: KS, YT, Methodology: KS, Investigation: KS, YT, Visualization: KS, Funding acquisition: KS, YT, Project administration: KS, YT, Supervision: KT, YT, Writing – original draft: KS, Writing – review & editing: KS, YT

## Declaration of Interests

The authors declare no competing interests.

## Notes

### Competing Interest Statement

The authors have declared no competing interest.

### Summary of Updates

We revised the Abstract and clarified in the Introduction the key problem addressed by this study.

