## Supplemental figures and table for "A naïve piRNA Surveillance System That Broadly Monitors the Germline Transcriptome for Adaptive Genome Defense"

Figure S1

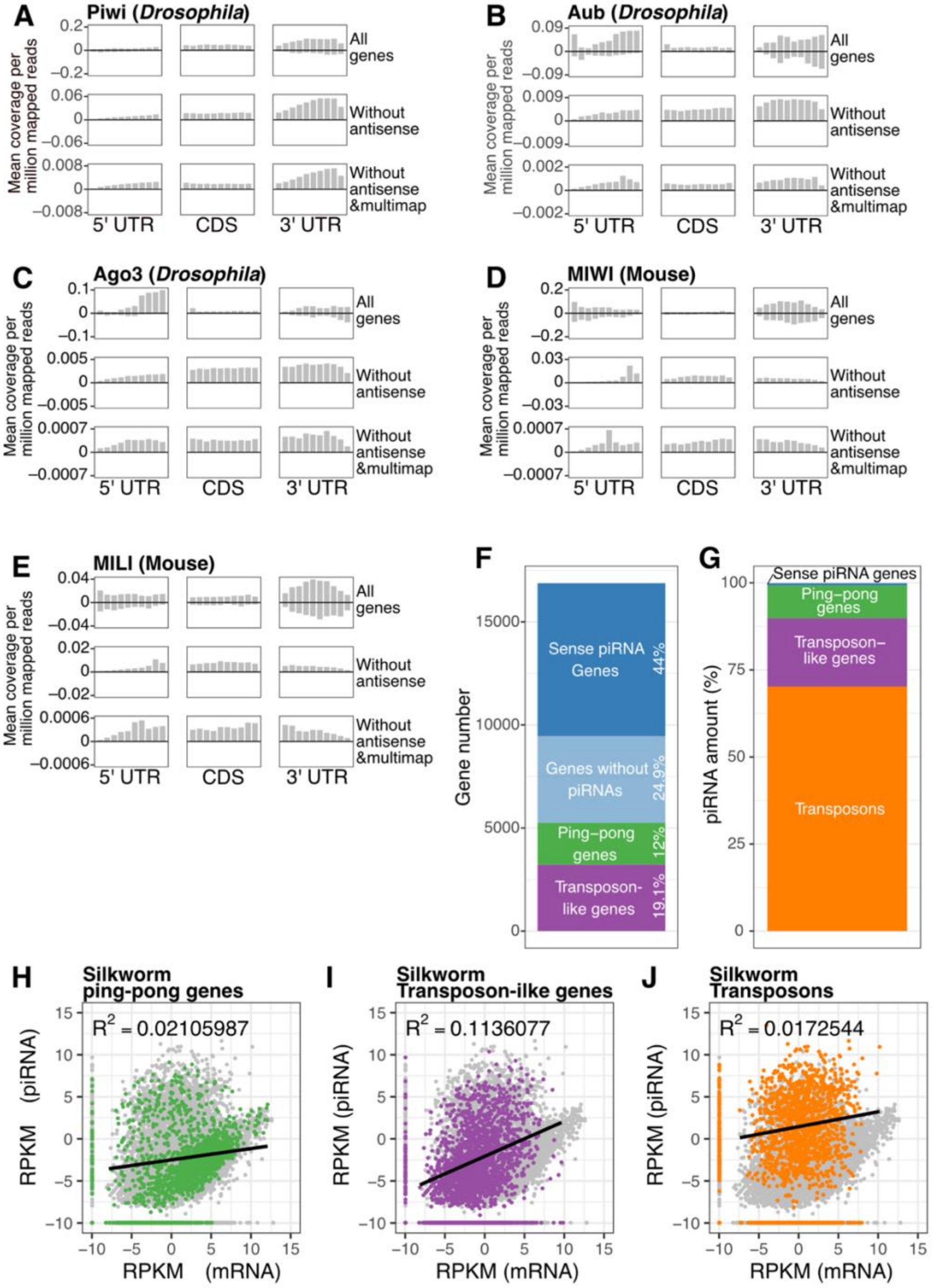


**Fig. S1. piRNAs derived from genes and transposons.**

(A–E) The distributions of the average piRNA coverage within 5' UTRs, CDSs and 3' UTRs of flies and mouse genes are shown. Results are shown for piRNAs bound to Piwi (A), Aub (B), and Ago3 (C) in flies, and MIWI (D) and MILI (E) in mice, respectively. For each gene, the UTRs and CDSs were each divided into 10 segments. The positive Y-axis indicates coverage in the sense direction, while the negative Y-axis indicates coverage in the antisense direction. Out of 30,570 *Drosophila* transcripts analyzed in total, 13,172 transcripts had no piRNAs mapped to antisense strand in three piRNA libraries. Out of 96,179 mice transcripts analyzed in total, 57,230 transcripts had no piRNAs mapped to antisense strand in two piRNA libraries.

(F) The percentage distribution of 16,880 genes within the four categories: sense piRNA genes, genes without piRNAs, ping-pong genes, and transposon genes. Genes without piRNAs refer to genes for which no piRNA reads were mapped in the total piRNA library.

(G) The percentage of piRNA abundance mapped to 16,880 genes and 1,811 transposons, with the total sum considered as 100%.

(H–J) Scatter plots illustrating the relationships between mRNAs and piRNAs (26–32 nt) mapped to the CDSs of 2,033 ping-pong genes (with at least one antisense piRNA read) (H), the CDSs of 3,220 transposon genes (homologous to transposons at the amino acid level) (I), and 1,811 different transposons (J) are shown on a $\log_2$ scale. A value of $\log_2(0.001)$ is added as the minimum value to both the vertical and horizontal axes. Regression lines were drawn using only those piRNAs and mRNAs whose levels were greater than $\log_2(0)$. The adjusted $R^2$ values are shown in the upper left of the graph.

### Figure S2

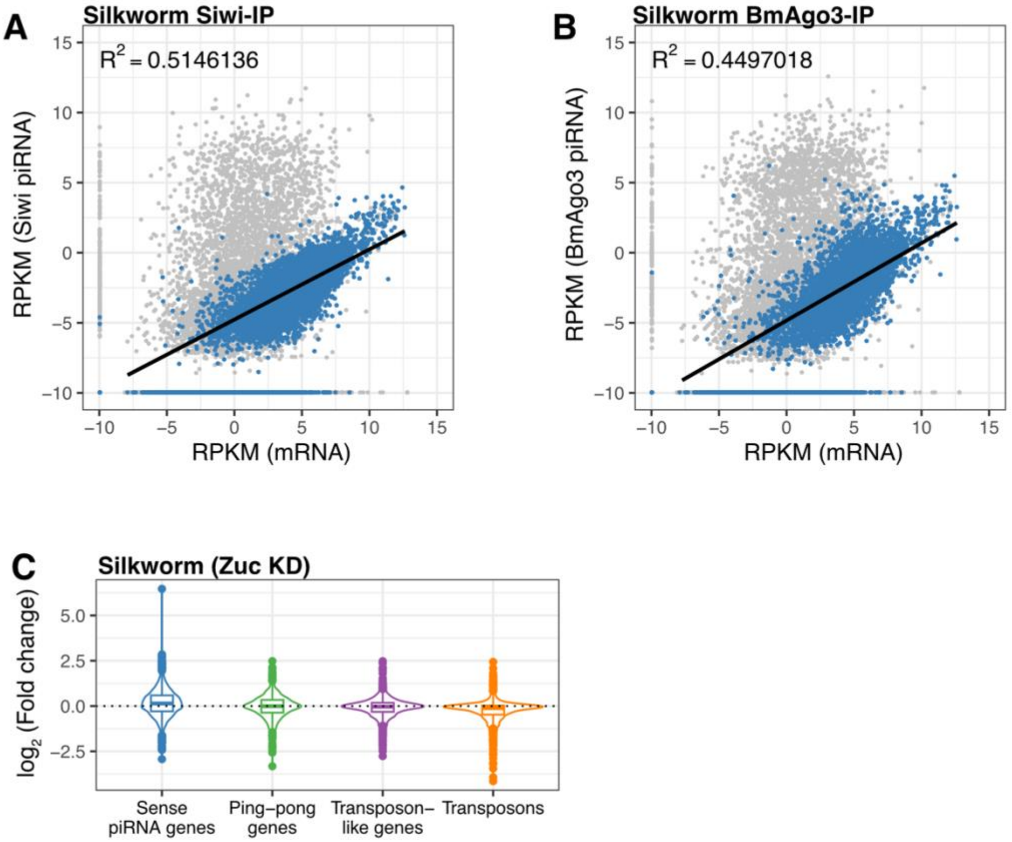


**Fig. S2. Gene-derived piRNAs are bona fide piRNAs.**

(A–B) Scatter plots illustrating the relationships between mRNAs and Siwi- (A) or BmAgo3-bound (B) piRNAs (26–32 nt) mapped to the CDSs of the genes on a $\log_2$ scale. A minimum value of $\log_2(0.001)$ is added to both the vertical and horizontal axes. The 7,422 sense piRNA genes are shown in blue. Regression lines were drawn using only those piRNAs and mRNAs whose levels were greater than $\log_2(0)$. The adjusted $R^2$ values are shown in the upper left of the graph.

(C) Violin plots of the change in piRNA abundance by Zuc KD. RPKM (Zuc KD)/RPKM (control KD) was calculated and shown for each category. Transcripts with an average RPKM greater than $\log_2(–2.5)$ were shown in these plots. Center line, median; box limits, upper and lower quartiles; whiskers, 1.5 × interquartile range; points, outliers. All points were shown in the inset. See also Figure 2B.

### Figure S3

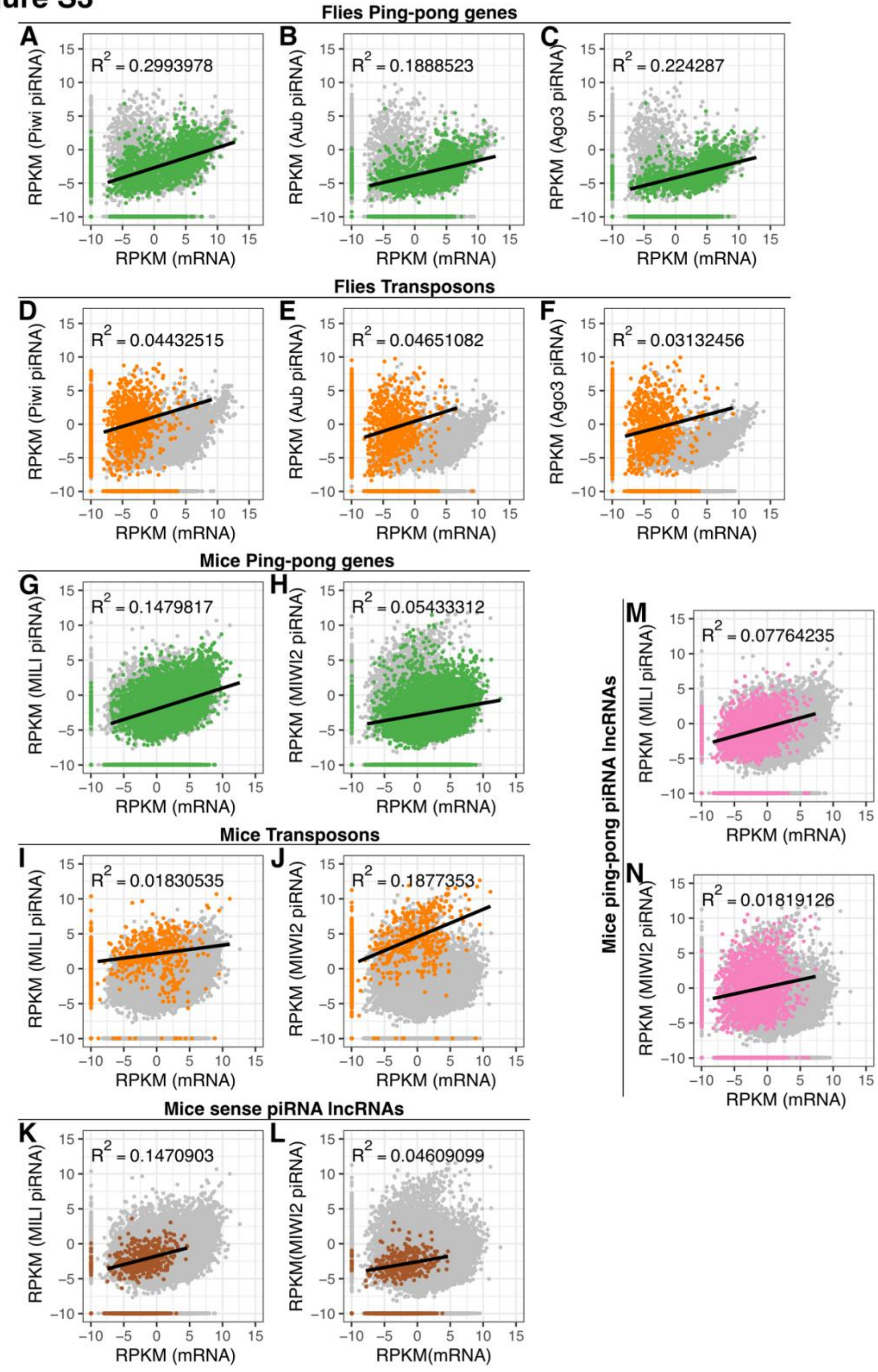


**Fig. S3. Gene-derived piRNAs in flies and mice.**

(A–F) The scatter plots show the relationship between ovarian mRNA abundance and PIWI-bound piRNAs in *Drosophila* on a $\log_2$ scale. Genes and transposons in each category were plotted: ping-pong piRNA genes with antisense piRNAs (A–C) and transposons (D–F). Results are shown for piRNAs bound to *Drosophila* Piwi (A, D), Aub (B, E), and Ago3 (C, F). Genes and transposons not belonging to these categories are shown in gray. A value of $\log_2(0.001)$ is added to both the vertical and horizontal axes as the minimum value. Regression lines were drawn using only those piRNAs and mRNAs whose levels were greater than $\log_2(0)$. The adjusted $R^2$ values are shown in the upper left of the graph.

(G–N) The scatter plots show the relationship between mRNA abundance and PIWI-bound piRNAs in mice on a $\log_2$ scale. Genes, lncRNAs, and transposons in each category were plotted: ping-pong piRNA genes with antisense piRNAs (G, H), transposons (I, J), lncRNAs without antisense piRNAs (K, L) and lncRNAs with antisense piRNAs (M, N). Results are shown for piRNAs bound to mouse MILI (G, I, K, M) and MIWI (H, J, L, N). A value of $\log_2(0.001)$ is added to both the vertical and horizontal axes as the minimum value. Regression lines were drawn using only those piRNAs and mRNAs whose levels were greater than $\log_2(0)$. The adjusted $R^2$ values are shown in the upper left of the graph.

### Figure S4

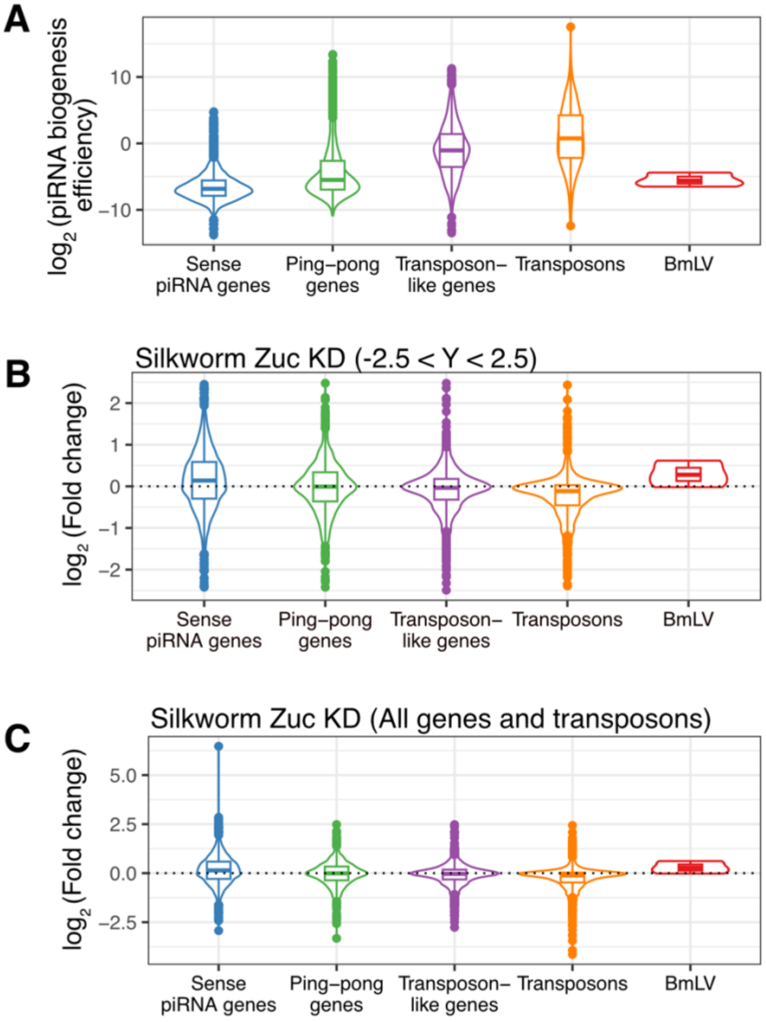


**Fig. S4. BmLV-derived piRNAs are produced by the same pathway as gene-derived piRNAs.**

(A) The violin plots show the piRNA biogenesis efficiency (piRNA amount/mRNA amount) for sense piRNA genes, ping-pong piRNA genes, transposon-like genes and transposons, and BmLV. See also Figure 1D. Center line, median; box limits, upper and lower quartiles; whiskers, 1.5 × interquartile range; points, outliers.

(B, C) Violin plots of the change in piRNA abundance by Zuc KD. RPKM (Zuc KD)/RPKM (control KD) was calculated and shown for each category. Transcripts with an average RPKM greater than $\log_2$(–2.5) were shown in these plots. Center line, median; box limits, upper and lower quartiles; whiskers, 1.5 × interquartile range; points, outliers. (B) Only those with Y-axis values between –2.5 and 2.5 were shown in the inset; 11 sense piRNA genes, 2 ping-pong genes, 1 transposon gene, and 10 transposons are excluded. See also Figure 2B. (C) All genes and transposons were shown.

### Figure S5

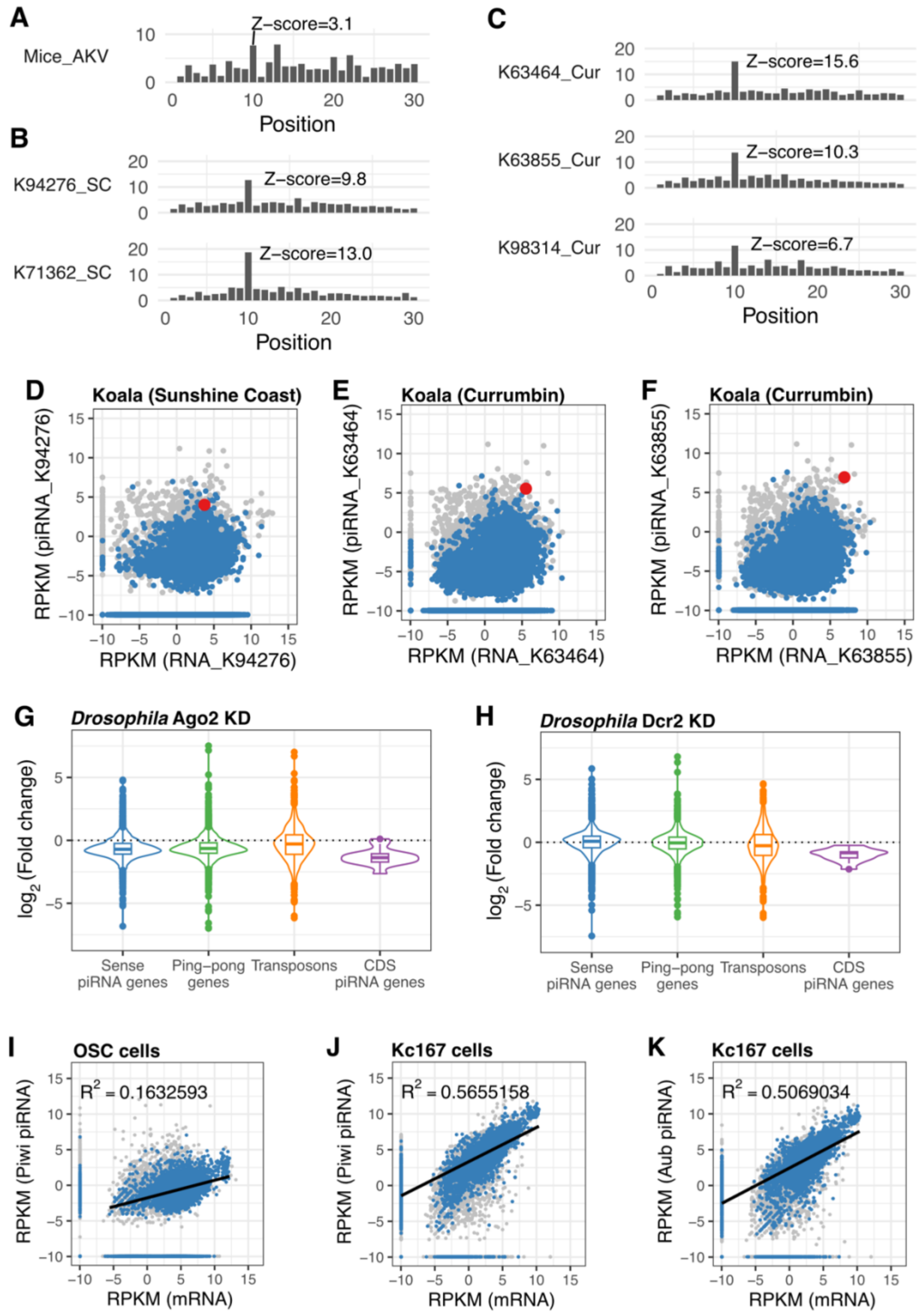


**Fig. S5. In vivo evidence for naïve piRNA biogenesis during retroviral endogenization.**

(A–C). Ping-pong signature analysis of retrovirus-derived piRNAs are shown. The distribution of 5′–5′ overlap lengths between sense and antisense piRNA pairs were computed (x-axis: overlap in nt, 0–30; y-axis: normalized pair %). The Z-score at 10 nt overlap is indicated in each panel. AKV: small-RNA reads from mouse testis (A). KoRV-A: small-RNA reads from koala testis—Sunshine Coast (B) and Currumbin (C).

(D–F) The scatter plots show the relationship between mRNA abundance and piRNAs (≥ 26 nt) in koala testes from Sunshine Coast (D), and Currumbin (E, F). In each sample, mRNAs and piRNAs were mapped to genes in the sense orientation. A total of 43,042 genes in the koala piRNA libraries that lacked antisense piRNA mapping are colored blue. KoRV-A is shown in red, and three endogenous retroviruses are shown in purple (E–F). A value of $\log_2(0.001)$ is added to both the vertical and horizontal axes as the minimum value.
(G, H) Violin plots of the change in piRNA abundance by *Drosophila Ago2* (G) or *Dicer* (H) KD for each category: sense piRNA genes, ping-pong piRNA genes, transposons and CDS piRNA genes (*37*). Center line, median; box limits, upper and lower quartiles; whiskers, 1.5 × interquartile range; points, outliers.
(I–K) The scatter plots show the relationship between mRNA abundance and Piwi-bound piRNAs in OSC cells (I) and Piwi- (J) or Aub (K)-bound piRNAs in Kc167 cells, all mapped to genes in the sense direction in *Drosophila melanogaster*. 7,106 genes with no piRNAs mapped to the antisense strand in all three fly libraries (Piwi, Aub, Ago3) used in Figure 3A–C are plotted. A value of $\log_2(0.001)$ is added to both the vertical and horizontal axes as the minimum value. Regression lines were drawn using only those piRNAs and mRNAs whose levels were greater than $\log_2(0)$. The adjusted $R^2$ values are shown in the upper left of the graph.

#### Table S1. Nucleotide sequences used in this study

miRNA sequence used for library nomalization

| Sequence name | Sequence |
|---|---|
| bmo-bantam | TGAGATCATTGTGAAAGCTAAT |
| bmo-let-7a | TGAGGTAGTAGGTTGTATAG |
| bmo-miR-1 | TGGAATGTAAAGAAGTATGGAG |
| bmo-miR-100 | AACCCGTAGATCCGAACTTGTG |
| bmo-miR-11 | CATCACAGTCAGAGTTCTAGCT |
| bmo-miR-184 | TGGACGGAGAACTGATAAGG |
| bmo-miR-2758 | ACTTGGTAGAACACGTAGTAAG |
| bmo-miR-2760* | CGAGGCTTAATTGAACCAAAAAGC |
| bmo-miR-277 | TAAATGCACTATCTGGTACGACA |
| bmo-miR-2778a | CAGAGTACGCAAAAAAACAATT |
| bmo-miR-278 | TCGGTGGGATCTTCGTCCGTTT |
| bmo-miR-279a | TGACTAGATCCACACTCATCCA |
| bmo-miR-279b | TGACTAGATCTACACTCATTGA |
| bmo-miR-317 | TGAACACAGCTGGTGGTATCTCAGT |

spike sequence used for library nomalization

| Sequence name | Sequence |
|---|---|
| spike1 | GTCCCACTCCGTAGATCTGTTC |
| spike2 | GATGTAACGAGTTGGAATGCAA |
| spike3 | TAGCATATCGAGCCTGAGAACA |
| spike4 | CATCGGTCGAACTTATGTGAAA |
| spike5 | GAAGCACATTCGCACATCATAT |
| spike6 | TCTTAACCCGGACCAGAAACTA |
| spike7 | AGGTTCCGGATAAGTAAGAGCC |
| spike8 | TAACTCCTTAAGCGAATCTCGC |
| spike9 | AAAGTAGCATCCGAAATACGGA |
| spike10 | TGATACGGATGTTATACGCAGC |

dsRNA preparation

| Sequence name | Sequence |
|---|---|
| dsRluc(*Renilla* luciferase)-F | TAATACGACTCACTATAGGGCCTTTCACTACTCCTACGAGC |
| dsRluc(*Renilla* luciferase)-R | TAATACGACTCACTATAGGGTGGAGCGTCCTCCTGGCTG |
| dsZuc-F | GCGTAATACGACTCACTATAGGCTAACTAGCACTGCCTACA |
| dsZuc-R | GCGTAATACGACTCACTATAGGCTCAAATTCAGTCTTAAACTG |
